# Network-based analysis of allele frequency distribution among multiple populations identifies adaptive genomic structural variants

**DOI:** 10.1101/2021.01.25.428140

**Authors:** Marie. Saitou, Naoki Masuda, Omer. Gokcumen

**Affiliations:** Dept. of Biological Sciences, University at Buffalo, State University of New York, Buffalo, NY 14260-2900, USA; Currently at the Faculty of Biosciences, Norwegian University of Life Sciences, Universitetstunet 3, 1430 Ås, Norway.; Dept. of Medicine, University of Chicago. Section of Genetic Medicine, 5841 S. Maryland Ave., Chicago, IL, 60637-1447, USA; Department of Mathematics, University at Buffalo, State University of New York, Buffalo, NY 14260-2900, USA; Computational and Data-Enabled Science and Engineering Program, University at Buffalo, State University of New York, Buffalo, NY 14260-5030, USA

**Author notes:** Correspondence: O.G., N.M.

## Abstract

Structural variants have a considerable impact on human genomic diversity. However, their evolutionary history remains mostly unexplored. Here, we developed a new method to identify potentially adaptive structural variants based on a network-based analysis that incorporates genotype frequency data from 26 populations simultaneously. Using this method, we analyzed 57,629 structural variants and identified 577 structural variants that show high population distribution. We further showed that 39 and 20 of these putatively adaptive structural variants overlap with coding sequences or are significantly associated with GWAS traits, respectively. Closer inspection of the haplotypic variation associated with these putatively adaptive and functional structural variants reveals deviations from neutral expectations due to (i) population differentiation of rapidly evolving multi-allelic variants, (ii) incomplete sweeps, and (iii) recent population-specific negative selection. Overall, our study provides new methodological insights, documents hundreds of putatively adaptive variants, and introduces evolutionary models that may better explain the complex evolution of structural variants.

## Introduction

Emerging technologies have recently revealed hundreds of thousands of genomic structural variants, including polymorphic duplications, deletions, inversions, and mobile transposable elements in the human genome (Feuk et al. 2006; Redon et al. 2006; Hurles et al. 2008; Conrad et al. 2010; Pang et al. 2010). Unlike single nucleotide variants, each structural variant affects a continuous block in the genome and thus is more likely to result in a phenotypic effect (Hurles et al. 2008; Weischenfeldt et al. 2013; Sudmant, Rausch, et al. 2015). Several structural variants have been documented to have considerable effects on human disease and evolution (Dennis and Eichler 2016; Payer et al. 2017; Hsieh et al. 2019; Ho et al. 2020; Saitou and Gokcumen 2020). Some of these functional variants reach >20% allele frequency in human populations, and some affect the copy number variation of entire protein-coding genes (McCarroll et al. 2005; Handsaker et al. 2015).

The poster child for adaptive structural variation in humans is the copy number variation of the amylase gene. Several studies put forward evidence for positive selection of higher amylase gene copy number in the human lineage, and further in high-starch-consuming human populations (Perry et al. 2007). Another striking example of potentially adaptive structural variants is the deletion of *LCE3B* and *LCE3C.* This variant is one of the leading susceptibility markers to psoriasis (de Cid et al. 2009). This deletion was shown to be retained in the human lineage since Human-Neanderthal divergence under balancing selection (Pajic et al. 2016), arguably maintaining a balance between protection against pathogens and facilitating immune-mediated disorders. Recently, a genome-wide analysis identified several Neanderthal- and Denisovan-introgressed structural variants that show strong signatures of adaptation in Oceanic populations (Hsieh et al. 2019). Collectively, these studies, along with others (see (Saitou and Gokcumen 2020) for a detailed review), imply that several common structural variants contribute to human phenotypic variation and may have evolved under diverse adaptive scenarios.

Despite the increasing appreciation of their role in human adaptive evolution, structural variants have not been scrutinized as much as single nucleotide variants due to technical difficulties. From a methodological perspective, structural variants are more challenging to discover and genotype due to their localization in highly repetitive sections of the genome. In addition, they are generated through complex mutational mechanisms, often involving gene conversions and unequal recombination events (Kidd et al. 2010; Handsaker et al. 2011; Lupski 2015; Carvalho and Lupski 2016; Sekar et al. 2016). As a result, structural variants are usually harbored by complex haplotypes, and they often are not tagged perfectly with flanking variants (Sudmant, Mallick, et al. 2015). The complexity of haplotypic architecture harboring structural variants complicates analyses of neutrality and integration of structural variants to genome-wide association studies. Most studies primarily interrogate single nucleotide variants and structural variants are often considered if only a “tag” single nucleotide variant can be found. For example, our work has resolved the complex evolutionary history of the common deletion of the metabolizing *GSTM1* gene. Locus-specific studies showed a strong association between this deletion and bladder cancer (The GSTM deletion is the risk allele, *p*=4×10^−11^) (Rothman et al. 2010). A haplotype-based analysis of the locus suggested that this deletion has formed multiple times through independent mutation events and undergone gene conversion events (Saitou, Satta, Gokcumen, et al. 2018). Thus, due to the lack of single nucleotide variants tagging this deletion, most genome-wide association studies and traditional selection scans did not include this deletion. Investigating individual haplotypes that harbor the deletion led us to identify one particular haplotype associated with the deletion that has been subject to a recent selective sweep in the East Asian populations (Saitou, Satta, and Gokcumen 2018).

A second factor that complicates the evolutionary study of structural variants is that some are multiallelic (Quinlan and Hall 2012; Handsaker et al. 2015). For example, the haptoglobin locus harbors two large multiallelic and recurrent structural variants that are not tagged by any single nucleotide variants. Only after careful, locus-specific resolution of haplotypic variation were they shown to be associated with cholesterol levels (Boettger et al. 2016). Similarly, *AMY1* (Perry et al. 2007), as we noted above, and *DMBT1 (Polley et al. 2015)* loci harbor multiallelic structural variations that were associated with dietary and metabolic traits.

However, even for amylase gene copy number variation, arguably the best-studied structural variant in the human genome from an evolutionary perspective, the timing and existence of putative adaptive forces remains elusive (Mathieson and Mathieson 2018). In fact, structural variants are often consciously left out from most selection scans along with segmental duplications and other repetitive regions due to the complications that we described above (**e.g.,**(Schrider and Kern 2017)). In sum, we argue that the full impact of structural variants on human evolution has not been understood and may explain some of the most exciting, yet to be described, adaptive variation in humans.

Given the complexity of haplotypes that harbor a considerable number of structural variants, measures that depend on accurate genotyping of haplotypic variation, such as allele frequency spectra (e.g., Tajima’s D (Tajima 1993)) or linkage-disequilibrium/homozygosity (e.g., iHS (Voight et al. 2006), XP-EHH (Sabeti et al. 2007)) are often underpowered. Instead, direct population differentiation metrics may be the most appropriate and unbiased way to identify putatively adaptive structural variants among human populations. Population-differentiation based methods are robust to haplotype disruption due to gene conversion, recurrence, or the presence of multiple alleles. Most studies that identify adaptive structural variants have employed population-differentiation based methods (Redon et al. 2006; Xue et al. 2008; Sudmant, Mallick, et al. 2015; Almarri et al. 2020; Bergström et al. 2020). Deviations from expected allele frequency distribution can provide information on several types of selection (positive, negative, or stabilizing), and differential selection with complex histories (selection on standing variation, recent geography-specific negative selection, oscillating selective forces such as dynamic environmental change (Key et al. 2014)). This is important because it has been shown that “classical” sweeps were rare (Hernandez et al. 2011) and selection on standing variants are likely to be the major force of human genomic adaptation (Schrider and Kern 2017), as recently shown for multiple alleles shaping skin color (Crawford et al. 2017; Martin et al. 2017).

To measure the population differentiation of genetic variants, F_ST_ statistics (Weir and Cockerham 1984) and V_ST_ statistics for copy number variants (Redon et al. 2006) are commonly used. More recent research has developed methods to compare multiple populations, primarily for admixture analysis (e.g., F_3_ statistics (Reich et al. 2009)). However, these methods can only compare two or three populations to each other. Recently, (Duforet-Frebourg et al. 2016) developed a PCA-based method to identify single nucleotide variants with population differentiation by analyzing 10 populations simultaneously. They confirmed well-known targets for positive selection *(EDAR, SLC24A5, SLC45A2, DARC)* and discovered new candidate genes *(APPBPP2, TP1A1, RTTN, KCNMA, MYO5C)*. Here, we developed a new, network-based method to identify adaptive structural variants with unusual allele frequency distribution with which one can analyze (1) multiallelic variants and (2) the distribution of genotype frequency in multiple populations collectively.

## Results and Discussion

### Network-Based Method to identify Structural Variants with High Population Differentiation

Inspired by the emerging work that integrates all available population differentiation information to understand demographic and adaptive trends (Duforet-Frebourg et al. 2016), we developed a new method based on the Bhattacharyya similarity metric specifically to identify putatively adaptive outliers among structural variants ((Bhattacharyya 1943)**, Methods, fig.1**).

**Figure 1.**
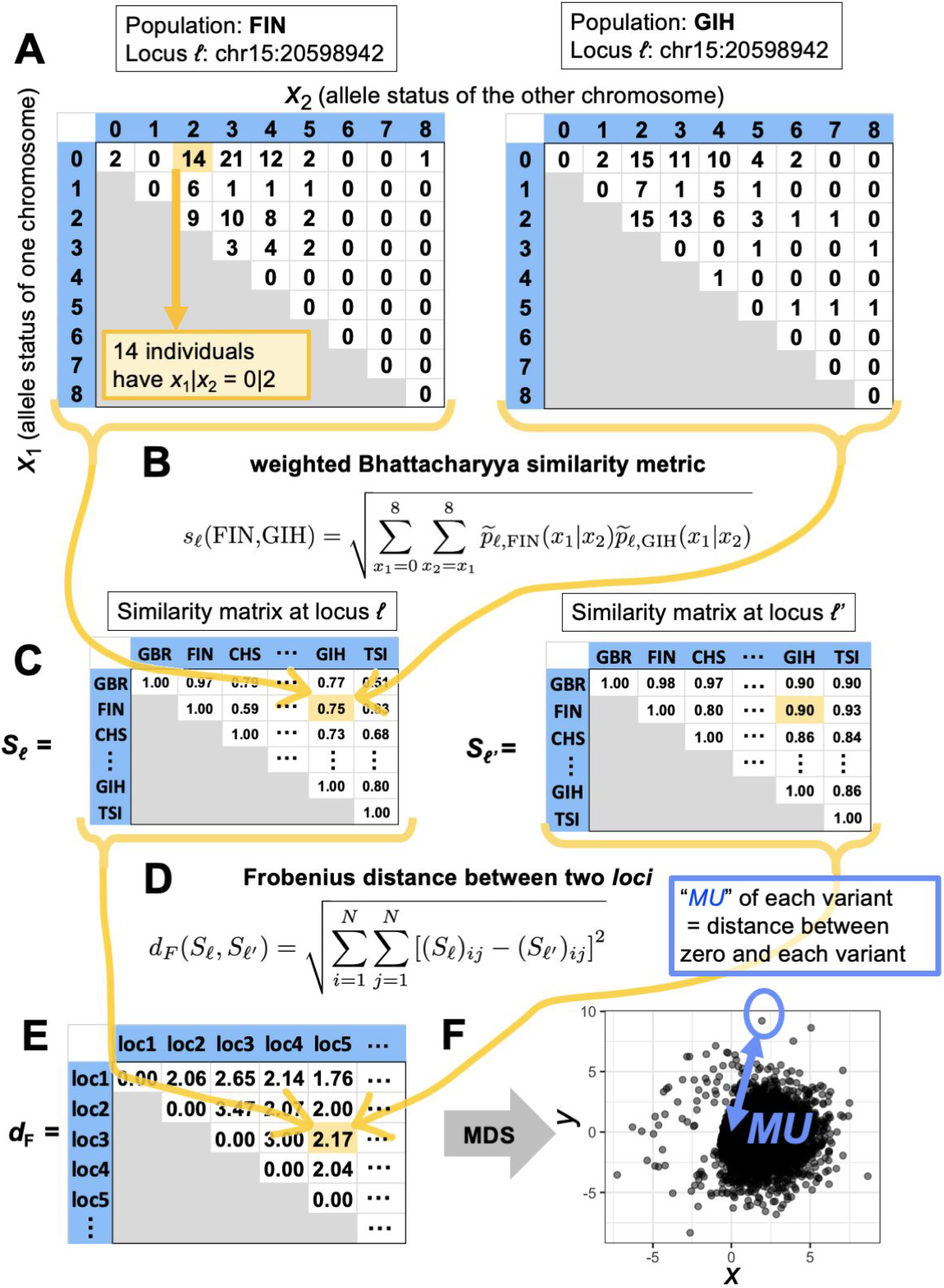
An overview of the calculation of “*Measure of Unusualness (MU)*”(see Methods for details). **A.** For each population and each locus, we have a 9 × 9 matrix representing the genotype. The row and column of the matrix represent one of the two chromosomes each. The cells contain the frequencies of specific genotype combinations. **B.** We calculate the similarity in the 9 × 9 matrices exemplified in panel A between each pair of populations. The similarity value ranges between 0 and 1. **C.** In this manner, for each locus, we obtain a 26 × 26 matrix representing the similarity between different pairs of populations. The diagonal entries of the similarity matrix are equal to 1 because any population is identical to itself, yielding the largest possible value of the similarity, which is 1. **D.** We calculate the distance between the 26 × 26 matrices for each pair of loci. **E.** In this manner; we obtain a distance matrix representing the distance between the different pairs of loci. **F.** We carry out the MDS to project the obtained distance matrix into the two-dimensional embedding space. Each circle represents a locus. The distance between the locus and the origin in the embedding space defines *MU* for each locus.

Briefly, we characterize each locus (*ℓ*) as an *N* ✕ *N* similarity matrix *S_ℓ_* based on the genotype frequency of the *N* = 26 populations in the 1000 Genomes Project phase 3 data set (Sudmant, Rausch, et al. 2015). We measure a modified Bhattacharyya similarity metric between each pair of populations based on the transformed probability distribution (for the original Bhattacharyya metric, see (Bhattacharyya 1943; Cha and Srihari 2002)). To increase the sensitivity to identify the population differentiation of variants with many alleles, we use a weighted variant of the Bhattacharyya similarity metric. Overall, we analyzed *M* = 58,644 variants, including 57,629 structural variants, 1,008 uniformly randomly chosen single nucleotide polymorphisms, as well as 7 single nucleotide polymorphisms that were reported to be under adaptive evolution (Norton et al. 2007; Mou et al. 2008; Kimura et al. 2009; Smith et al. 2009; Basu Mallick et al. 2013; Ding et al. 2013; Ko et al. 2013; Wilde et al. 2014; Wu et al. 2016; Deng and Xu 2018). We constructed a distance matrix for each of the *M* loci and compared the similarity matrix *S_ℓ_* across all these loci (*ℓ* = 1, …, *M*). We define the distance between *S_ℓ_* (at one locus) and *S_ℓ′_* (at another locus) by the Frobenius norm, denoted by *d*_F_. The *M* ✕ *M* Frobenius distance matrix, denoted by *F*, tabulates the difference between each pair of loci, and its (*ℓ*, *ℓ′*) entry is given by *d*_F_(*S_ℓ_*, *S_ℓ_*_′_). These steps provided us a matrix indicating how structural variants relate to each other based on their global genotype frequency distribution.

We assumed, based on previous literature (Conrad et al. 2010), that the majority of structural variants will be evolving under neutrality or near neutrality. Therefore, population differentiation should primarily be driven by genetic drift. We then reasoned that structural variants that have unusual allele frequency distribution among the 26 populations as compared to the genome-wide observations are likely to have evolved under non-neutral conditions. To visualize the relationships between structural variants based on their global allele frequency distribution, we ran a multidimensional scaling (MDS) algorithm. To empirically measure these relationships, we calculated the distance between the origin and each variant in the MDS space and defined it as “*Measure of Unusualness (MU)*”, or the degree of unusual allele frequency distribution (**Methods, fig. 1**). This measure informs on the unusualness of global population differentiation of a given structural variant, as compared to the entirety of the data set.

### Empirical Confirmation

In addition to the structural variants, we also included 1,008 random single nucleotide polymorphisms (SNPs), six SNPs which are known for high population differentiation, and one SNP under balancing selection in our pipeline. (**fig. 2A, TableS1, Methods**). We reasoned that the uniformly random control SNPs will provide an additional marker set whereby differentiation is determined by near-neutral forces. Using this data set, we confirmed that the distribution of *MU* measured for structural variants is not significantly different from that measured for the uniformly randomly chosen SNPs from the 1000 Genome Phase 3 data set (Sudmant, Rausch, et al. 2015) (*p* = 0.88, Mann-Whitney test, **fig. S1**), suggesting that similar to single nucleotide variants, overall distribution of *MU* for SVs is neutral-like.

**Figure 2.**
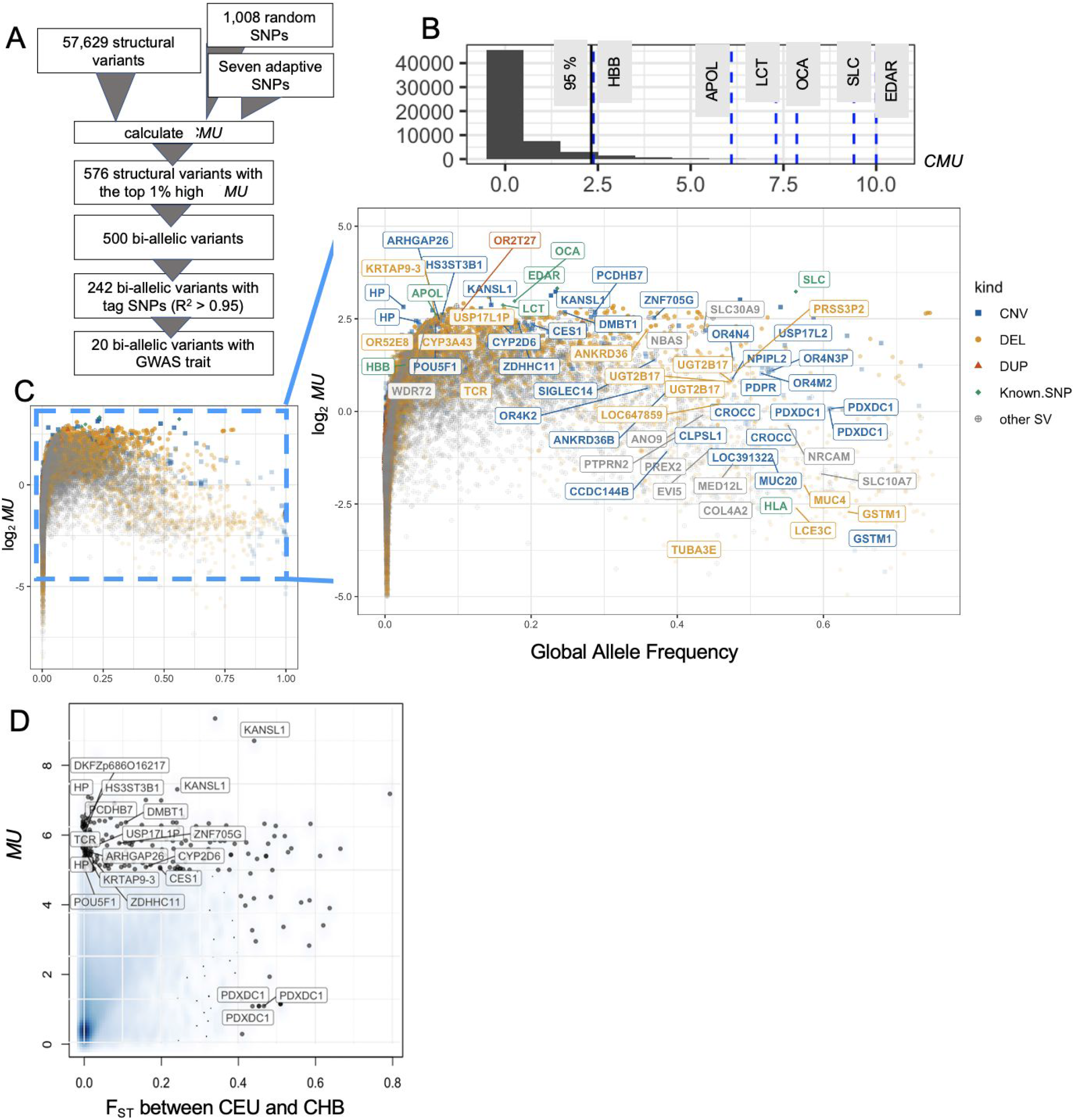
An overview of the study pipeline and results. **A**. The bioinformatic pipeline for the calculation of *MU* (see **Methods** for details). **B**. The histogram of *MU* of the 1,008 randomly selected SNPs and the six “adaptive SNPs” that were found to be under selection in previous studies (see **Methods**). The latter group was indicated by blue vertical dashed lines on the histogram, and the genes affected by these variants were labeled. The 95^th^ percentile of the distribution was marked by a black vertical line. **C.** The relationship between allele frequency and *MU*. The horizontal axis indicates the global alternative allele frequency. The vertical axis indicates the logarithm of the *MU* value. The exonic structural variants with *MU* > 5 or global allele frequency > 0.25 are labelled, as well as the six “adaptive SNPs” and HLA SNP. Some gene names are shown multiple times (e.g., HP, KANSP1, and PDXDC1); this happens because multiple structural variants overlapping these genes were reported in the 1000 Genomes Project Phase 3 data set. Colors represent different types of variants. The abbreviation is from the 1000 Genome Project phase 3 structural variants data set., CNV; copy number variants (multiallelic variants), DEL: deletion, DUL: duplication, Known. SNP: SNPs from previous studies (see Methods), other SVs (insertion, inversion, Alu, Long interspersed nuclear element, SINE-VNTR retrotransposons). **D.** Comparison of F_ST_ (Weir and Cockerham 1984) between CEU and CHB populations and *MU*. Bi-allelic structural variants with F_ST_ (between CEU and CHB) >0.4 and *MU* > 5 were labeled. The shade in blue represents the density of the structural variants.

Next, we investigated the sensitivity of our method by measuring *MU* from specific SNPs that were previously shown to be evolving under local adaptive forces, including rs3827760 in *EDAR*, rs1426654 in *SLC24A5*, rs12913832 in *OCA*, rs4988235 in *LCT*, rs73885319 in *APOL*, and rs334 in *HBB* (**Methods**). These SNPs are well-studied and repeatedly reported likely to be under population-specific adaptations, and how they may functionally contribute to fitness in certain populations has been analyzed (**Methods**). Thus, they provide an appropriate gold standard to test the sensitivity of our method. We found that all of these six SNPs were shown to be in the top 5% of the *MU* distribution, improving our confidence in our method (**fig. 2B**). Even though MU was not designed to test balancing selection we also included in our analysis the rs1129740, which resides in the *HLA* locus and has been one of the handfuls of variants in humans that are thought to have evolved under long-term balancing selection (Teixeira et al. 2015). This allowed us to observe how *MU* behaves for such variants. We found that this non-synonymous mutation shows low *MU* values when considering its allele frequency (**fig. 2C**). We found it noteworthy that a structural variant, *LCE3BC* gene deletion, that we speculated previously to have evolved under balancing selection (Lin et al. 2015; Pajic et al. 2016; Saitou, Satta, and Gokcumen 2018) show similarly low *MU* values despite their high allele frequency (**fig. 2C**). Thus, it is plausible that high allele frequency structural variants that may have been evolving under balancing selection may exhibit unusually low *MU* values (**table S2**).

To understand the differences between more traditional methods of measuring population differentiation and our method, we compared our *MU* with direct allele-frequency-based F_ST_ between representative continental populations (**fig. 2D, table S3**). As expected, we found a significant correlation between these two measures (Spearman rank correlation coefficient > 0.49 and *p* < 10^−15^ for all comparisons). However, we also found notable discrepancies. We noted 125 variants that are in the top 1^ST^ percentile for *MU*, but show small F_ST_ (F_ST_ < 0.2 in any pairwise comparison of European (CEU), East Asian (CHB), and African (YRI) populations), including deletions that overlap with *MYOM1* (Myomesin-1) and *HS3ST3B1* (Heparan Sulfate-Glucosamine 3-Sulfotransferase 3B1) genes (**Table S3**). Closer inspection of these variants suggests that *MU* captures multi-allelic variations (**fig. 3A**) and within-continent variation (**fig. 3B**), which fell below the detection threshold of standard pairwise population comparisons. In addition, we noted structural variants that have large values of F_ST_ but do not stand out in terms of *MU*. For example, esv3643467 shows high among-continent population allele frequency differentiation (F_ST (CEU vs. CHB)_ = 0.41), but with relatively low *MU* values (*MU* = 0.24) (**fig. 3C, fig. S2**), likely due to serial founder effects that define human genetic variation as described previously (Ramachandran et al. 2005). That suggests that a gradual population differentiation may not lead to a high *MU* value in our measure. This is an advantage of our method because the ecological and cultural variation *within* rather than *among* continents often underlie adaptive evolution in humans. Examples include high altitude adaptation in Tibet (Beall et al. 2010) and Ethiopia (Scheinfeldt et al. 2012), the selection on height in African and American rainforests (Lopez et al. 2019), dietary adaptations within all continents (Sabeti et al. 2006; Mathieson and Mathieson 2018), and adaptation against malaria in Africa (Leffler et al. 2017) and in Asia (McManus et al. 2017). Thus, we argue that variants that have high among-continental differences that the standard F_ST_ detects often reflect the effects of major bottleneck/drift events (Ochoa and Storey 2019), such as out-of-Africa migrations, rather than adaptive sweeps. By contrast, our method, *MU*, which compares allele frequency distributions of 26 populations simultaneously, does not identify such variations as unusual. Overall, our *MU* method shows promise in capturing hundreds of novel putatively adaptive variants that have not been captured by traditional pairwise population comparisons.

**Figure 3.**
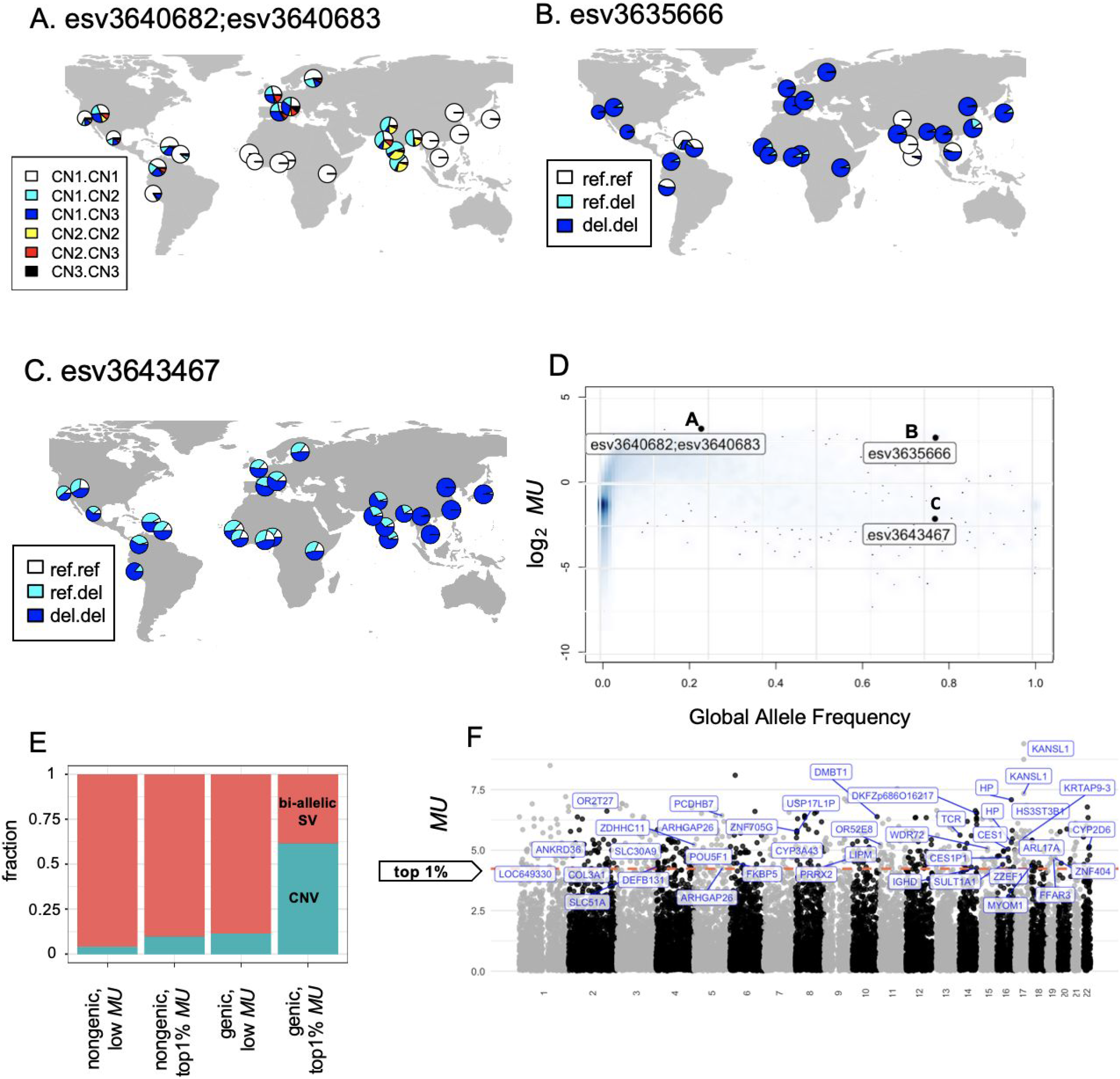
Examples of unusually distributed variants. **ABC** The geographical distribution of several identified structural variants as examples. Color represents genotype. White refers to a homozygous reference genotype. Non-white colors represent other genotypes. **A**. A multiallelic copy number variant; esv3640682;esv3640683, a multiallelic structural variant with unusual population differentiation and a top 1% *MU* value. CN means copy number, for example. CN1 is the copy number one allele and CN1.CN1 is the homozygous copy number one genotype. **B**. esv3635666 with unusual population differentiation and a top 1% *MU* value; this variant overlaps with the coding sequence of DKFZp686O16217 gene. **C**. esv3364367 at MUC4 exon is very common among human populations but showed low *MU* (also see **D**). **D**. The global allele frequency and *MU* of the highlighted variants (black points). The shade in blue represents the density of all the structural variants. **E** The relative ratio of bi-allelic structural variants and multiallelic copy number variants in each grouping based on exonic overlap (structural variants overlap with exonic sequences vs. those that do not) and *MU* values (top 1% vs. the rest). Bi-allelic SV: biallelic structural variants. CNV: multiallelic copy number variants with three or more alleles. **F.** Manhattan plots of *MU* of structural variants. The horizontal axis shows the chromosomal location of structural variants, and the vertical axis shows the *MU* value. The exonic variants with the top 1% *MU* (higher than 4.23) were labeled.

### *MU* Identifies Dozens of Putatively Adaptive Multiallelic and Exonic Structural Variants That Are Invisible to Traditional Selection Scans and Association Studies

There are several outstanding questions concerning the enrichment of specific properties of adaptive structural variants, including their functional relevance, the mutation mechanisms through which the variants are generated, and their size distribution. However, there are tremendous technical biases inherent in short-read sequencing based characterization of these variants, especially concerning extremely high false-negative rates in the discovery of certain types of structural variants, such as tandem and dispersed duplications and inversions (Kronenberg et al. 2015). Thus, instead of searching for general trends in our data set (e.g., adaptively evolving structural variants are larger or smaller than neutrally evolving ones), we focused on resolving the evolutionary forces shaping individual structural variants with functional implications.

In this spirit, we first investigated structural variants that overlap with coding sequences. We identified 39 structural variants with the top 1% *MU* value that contain one or more entire exon (**Table S4**). Many of these exonic structural variants were associated with metabolic traits and diseases in previous locus-specific analyses and include members of cytochrome p450 (*CYP3A43*, *CYP2D6*), solute carrier (*SLC30A9*, *SLC51A*), olfactory receptor (OR2T27, OR52E8) gene families. For example, *DMBT1* gene copy number was noted for its population differentiation and associated with dietary subsistence strategies (Polley et al. 2015). Similarly, the copy number variation affecting the *CES1 (Zhu and Markowitz 2013), CYP2D6 (Candiotti et al. 2005), HS3ST3B1 (Kim et al. 2010)*, and *SULT1 (Hebbring et al. 2008)* are associated with differences in metabolizing of xenobiotic substances, primarily described within a pharmacogenomics context. Interestingly, we found that 24 (~62%) of these exonic structural variants are multiallelic (**fig. 3E, 3F),** more than five times higher than genome-wide expectations (p = 0.0005, Chi-square test).

Strikingly, the exonic structural variants with the highest *MU* have been invisible to previous genome-wide association studies and selection scans. We argue that this is primarily because the current GWAS pipelines interrogate single nucleotide variants. Single nucleotide variants may not tag multiallelic structural variants due to gene conversion, recurrence, and potential genotyping errors, as discussed in the introduction. This phenomenon was diligently dissected by Boettger et al. (Boettger et al. 2016) for the haptoglobin (HP) locus, which harbors two recurrent and multiallelic exonic copy number variants that we found to show unusually high *MU*. They described a novel way to use a combination of single nucleotide variants to impute these structural variants in the locus. Their reanalysis of the genome-wide association studies revealed a previously hidden association between copy number variation affecting HP gene function and blood cholesterol levels. Based on our results, we argue that the effects of multiallelic structural variants on human evolution and phenotypic variation remain underappreciated.

### Resolving the Haplotypes of Putatively Adaptive Structural Variants to Investigate Their Evolutionary and Functional Effects

The complex evolutionary dynamics of structural variants often do not fit classical population genetics expectations, such as complete classical sweeps. Thus, we argue that careful investigation of the evolutionary histories of a few examples can provide valuable insights that can later be generalized at the genome-wide scale. Therefore, we wanted to resolve the haplotypes that harbor structural variants in order to investigate functional associations, coalescence times, and signatures of selection concerning these variants in more detail. There are 344 bi-allelic structural variants that are in the 1st percentile in terms of *MU* (> 4.23) and have strong linkage disequilibrium with nearby single nucleotide variants (R^2^ > 0.95) (**fiig. 2A, Table S4**)

Among these, we identified 20 haplotypes that are significantly associated with phenotypes (nominal p < 10^−9^; GWAS Atlas (https://atlas.ctglab.nl/; last accessed 3.23.2020) (**Table 2**). The selected 20 loci provided us with a means to further investigate the evolutionary and functional effects of structural variants that show unusual geographical distribution.

**Table 1.** Twenty-five multiallelic structural variants (CNV, copy number variation in **fig. 2C**) with the top 1% *MU* (higher than 4.23). Variant information is retrieved from the 1000 Genomes Project phase 3 data set (Sudmant, Rausch, et al. 2015). Start and End refer to the starting and ending locations of variants on the chromosome, respectively. We described the gene name if the structural variant contains one or more entire exon(s) of UCSC Genes.

| ID | Number of Alleles | MU | Chromosome | Start | End | Allele frequency | Gene name |
| --- | --- | --- | --- | --- | --- | --- | --- |
| esv3640682;esv3640683 | 3 | 9.40 | 17 | 44341412 | 44366497 | 0.234 | NA |
| esv3640680;esv3640681 | 3 | 8.75 | 17 | 44230893 | 44262697 | 0.227 | KANSL1 |
| esv3608414;esv3608415 | 3 | 8.09 | 6 | 26739614 | 26748928 | 0.486 | NA |
| esv3640677;esv3640678 | 3 | 7.34 | 17 | 44165338 | 44211686 | 0.145 | KANSL1 |
| esv3638992;esv3638993;esv3638994;esv3638995;esv3638996;esv3638997 | 7 | 7.08 | 16 | 72094527 | 72110961 | 0.025 | HP |
| esv3616125;esv3616126;esv3616127;esv3616128 | 5 | 7.03 | 8 | 8072609 | 8077628 | 0.541 | NA |
| esv3596068;esv3596069 | 3 | 6.85 | 3 | 46803577 | 46851159 | 0.131 | NA |
| esv3589567;esv3589568 | 3 | 6.80 | 1 | 248867276 | 248876082 | 0.089 | NA |
| esv3631446;esv3631447 | 3 | 6.64 | 13 | 21537857 | 21541131 | 0.065 | NA |
| esv3606964;esv3606965;esv3606966;esv3606967;esv3606968;esv3606969;esv3606970 | 8 | 6.43 | 5 | 140554408 | 140558942 | 0.288 | PCDHB7 |
| esv3624777;esv3624778;esv3624779;esv3624780 | 5 | 6.38 | 10 | 124344431 | 124353237 | 0.246 | DMBT1 |
| esv3640025;esv3640026 | 3 | 6.11 | 17 | 14224374 | 14483419 | 0.078 | HS3ST3B1 |
| esv3635703;esv3635704;esv3635705;esv3635706 | 5 | 6.06 | 14 | 107147166 | 107164532 | 0.287 | NA |
| esv3647555;esv3647556;esv3647557;esv3647558 | 5 | 6.01 | 22 | 30456542 | 30462376 | 0.079 | NA |
| esv3626640;esv3626641;esv3626642;esv3626643;esv3626644 | 6 | 5.87 | 11 | 63198026 | 63202706 | 0.282 | NA |
| esv3646343;esv3646344;esv3646345 | 4 | 5.82 | 20 | 62949512 | 62957862 | 0.197 | NA |
| esv3616121;esv3616122;esv3616123;esv3616124 | 5 | 5.80 | 8 | 7774146 | 7782327 | 0.586 | NA |
| esv3628214;esv3628215;esv3628216 | 4 | 5.80 | 12 | 147081 | 151800 | 0.173 | NA |
| esv3616116;esv3616117;esv3616118;esv3616119 | 5 | 5.78 | 8 | 7212582 | 7227421 | 0.368 | ZNF705G |
| esv3589783;esv3589784 | 3 | 5.77 | 2 | 9925419 | 9928927 | 0.158 | NA |
| esv3607012;esv3607013 | 3 | 5.56 | 5 | 142263109 | 142447062 | 0.074 | ARHGAP26 |
| esv3647175;esv3647176;esv3647177;esv3647178 | 5 | 5.54 | 22 | 16050654 | 16063474 | 0.143 | NA |
| esv3631107;esv3631108 | 3 | 5.45 | 12 | 127817403 | 128009847 | 0.079 | NA |
| esv3600641;esv3600642;esv3600643;esv3600644 | 5 | 5.45 | 4 | 60324149 | 60330809 | 0.043 | NA |
| esv3611756;esv3611757 | 3 | 5.43 | 7 | 151255 | 160130 | 0.118 | NA |

**Table 2.**
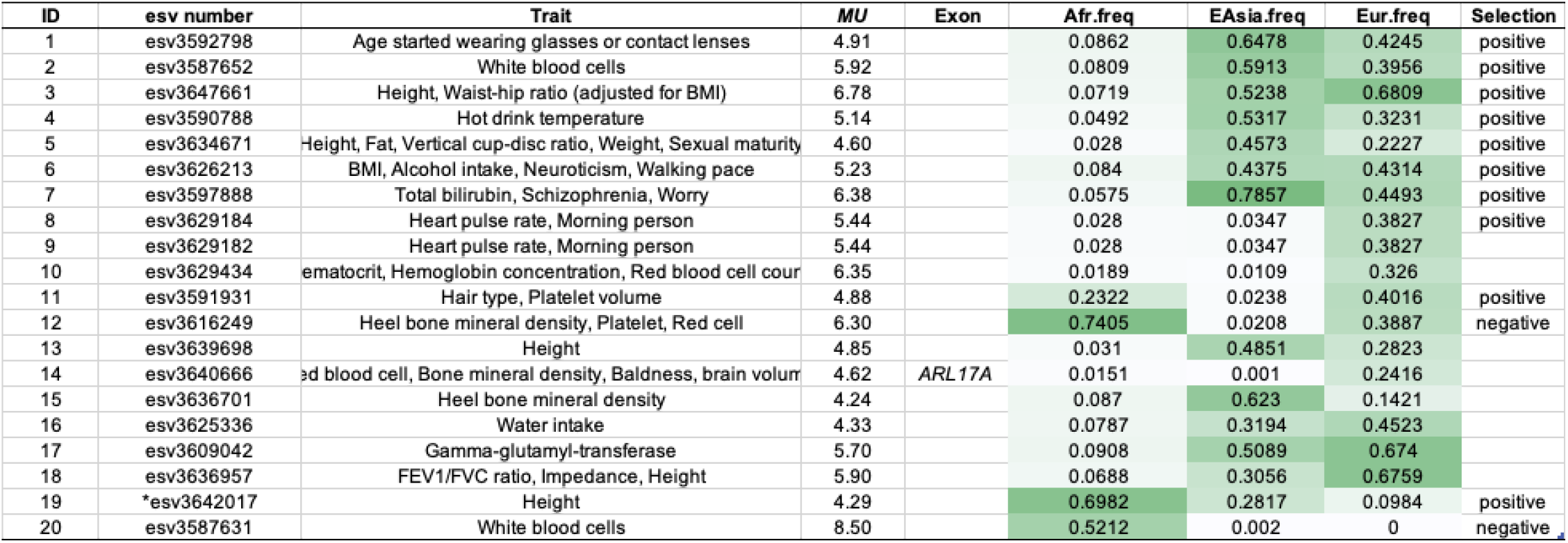
Structural variants with the top 1 % *MU* value and observed phenotypic effects through tag SNP (R^2^ > 0.95) in GWAS Atlas. All the structural variants in the table are deletions. ID is the same as **fig. 4**. We described the GWAS traits in the “Trait” column if the structural variant shows phenotypic effects through tag SNPs on GWAS Atlas. The linkage disequilibrium information for these structural variants and tag SNPs is found in **Table S5**. We described the gene name in the “Exon” column if the structural variant contains one or more entire exons of UCSC Genes. “Afr/EAsia/Eur.freq” are the frequency of the alternative allele in each population, Africa, East Asia and Europe. “Selection” column describes estimated natural selection based on the neutrality tests (**fig. 4**). Specifically, “negative” indicates cases where we found recent selection favoring the ancestral allele, while “positive” indicates cases where we found recent selection favoring the derived alleles. *This “deletion (esv3642017)” found in the 1000 Genomes samples as compared to the reference genome is actually a derived insertion that happens to be represented in the reference genome (Anagnou et al. 1988; Conrad et al. 2010; Schrider et al. 2013). Thus, even though the putative action of selection is on the non-deleted haplotypes, given that this haplotype carries the derived allele, we categorized the selection as “positive” in this case.

| ID | esv number | Trait | *MU* | Exon | Afr.freq | EAsia.freq | Eur.freq | Selection |
| --- | --- | --- | --- | --- | --- | --- | --- | --- |
| 1 | esv3592798 | Age started wearing glasses or contact lenses | 4.91 |  | 0.0862 | 0.6478 | 0.4245 | positive |
| 2 | esv3587652 | White blood cells | 5.92 |  | 0.0809 | 0.5913 | 0.3956 | positive |
| 3 | esv3647661 | Height, Waist-hip ratio (adjusted for BMI) | 6.78 |  | 0.0719 | 0.5238 | 0.6809 | positive |
| 4 | esv3590788 | Hot drink temperature | 5.14 |  | 0.0492 | 0.5317 | 0.3231 | positive |
| 5 | esv3634671 | Height, Fat, Vertical cup-disc ratio, Weight, Sexual maturity | 4.60 |  | 0.028 | 0.4573 | 0.2227 | positive |
| 6 | esv3626213 | BMI, Alcohol intake, Neuroticism, Walking pace | 5.23 |  | 0.084 | 0.4375 | 0.4314 | positive |
| 7 | esv3597888 | Total bilirubin, Schizophrenia, Worry | 6.38 |  | 0.0575 | 0.7857 | 0.4493 | positive |
| 8 | esv3629184 | Heart pulse rate, Morning person | 5.44 |  | 0.028 | 0.0347 | 0.3827 | positive |
| 9 | esv3629182 | Heart pulse rate, Morning person | 5.44 |  | 0.028 | 0.0347 | 0.3827 |  |
| 10 | esv3629434 | Hematocrit, Hemoglobin concentration, Red blood cell count | 6.35 |  | 0.0189 | 0.0109 | 0.326 |  |
| 11 | esv3591931 | Hair type, Platelet volume | 4.88 |  | 0.2322 | 0.0238 | 0.4016 | positive |
| 12 | esv3616249 | Heel bone mineral density, Platelet, Red cell | 6.30 |  | 0.7405 | 0.0208 | 0.3887 | negative |
| 13 | esv3639698 | Height | 4.85 |  | 0.031 | 0.4851 | 0.2823 |  |
| 14 | esv3640666 | Red blood cell, Bone mineral density, Baldness, brain volume | 4.62 | ARL17A | 0.0151 | 0.001 | 0.2416 |  |
| 15 | esv3636701 | Heel bone mineral density | 4.24 |  | 0.087 | 0.623 | 0.1421 |  |
| 16 | esv3625336 | Water intake | 4.33 |  | 0.0787 | 0.3194 | 0.4523 |  |
| 17 | esv3609042 | Gamma-glutamyl-transferase | 5.70 |  | 0.0908 | 0.5089 | 0.674 |  |
| 18 | esv3636957 | FEV1/FVC ratio, Impedance, Height | 5.90 |  | 0.0688 | 0.3056 | 0.6759 |  |
| 19 | *esv3642017 | Height | 4.29 |  | 0.6982 | 0.2817 | 0.0984 | positive |
| 20 | esv3587631 | White blood cells | 8.50 |  | 0.5212 | 0.002 | 0 | negative |

Using the linked haplotypic variation for the 20 structural variants, we retrieved allele ages from the Human Genome Dating database (Albers and McVean 2020) (https://human.genome.dating; last accessed, 3.23.2020). Under neutrality, an allele’s age is expected to positively correlate with its allele frequency (Patterson 2005). Given that we are explicitly investigating variants that are putatively evolving under population-specific adaptive forces, we expect deviations from this expectation. **Figure 4A-C** shows the estimated age of the allele and its frequency in European, East Asian, and African populations, respectively. If a variant has emerged recently, but its frequency is common (left upper side) in a given population, it suggests a potential recent selective sweep (i.e., a new allele is rapidly favored and increases its frequency). In contrast, if a variant is old and its frequency is rare, these are candidates for recent negative selection against the allele in that particular population. To more formally interrogate this line of inquiry, we calculated how long it takes for a new allele to reach at a given frequency in each population under neutrality using formula (15) in (Kimura and Ohta 1973) assuming previously published demographic parameters (Schaffner et al. 2005) (**fig. S3**). In a manner similar to allele frequency expectations, the age estimate of a variant older than the neutral estimation may suggest a faster increase in allele frequency and a recent selective sweep (**fig. 4A-4C**). In parallel, we calculated Tajima’s D scores of 5k upstream and downstream regions of the 20 structural variants of interest and the iHS scores of the tag single nucleotide variants of the target structural variants (**Methods**). We summarized these values in **fig. 4D**.

**Figure 4:**
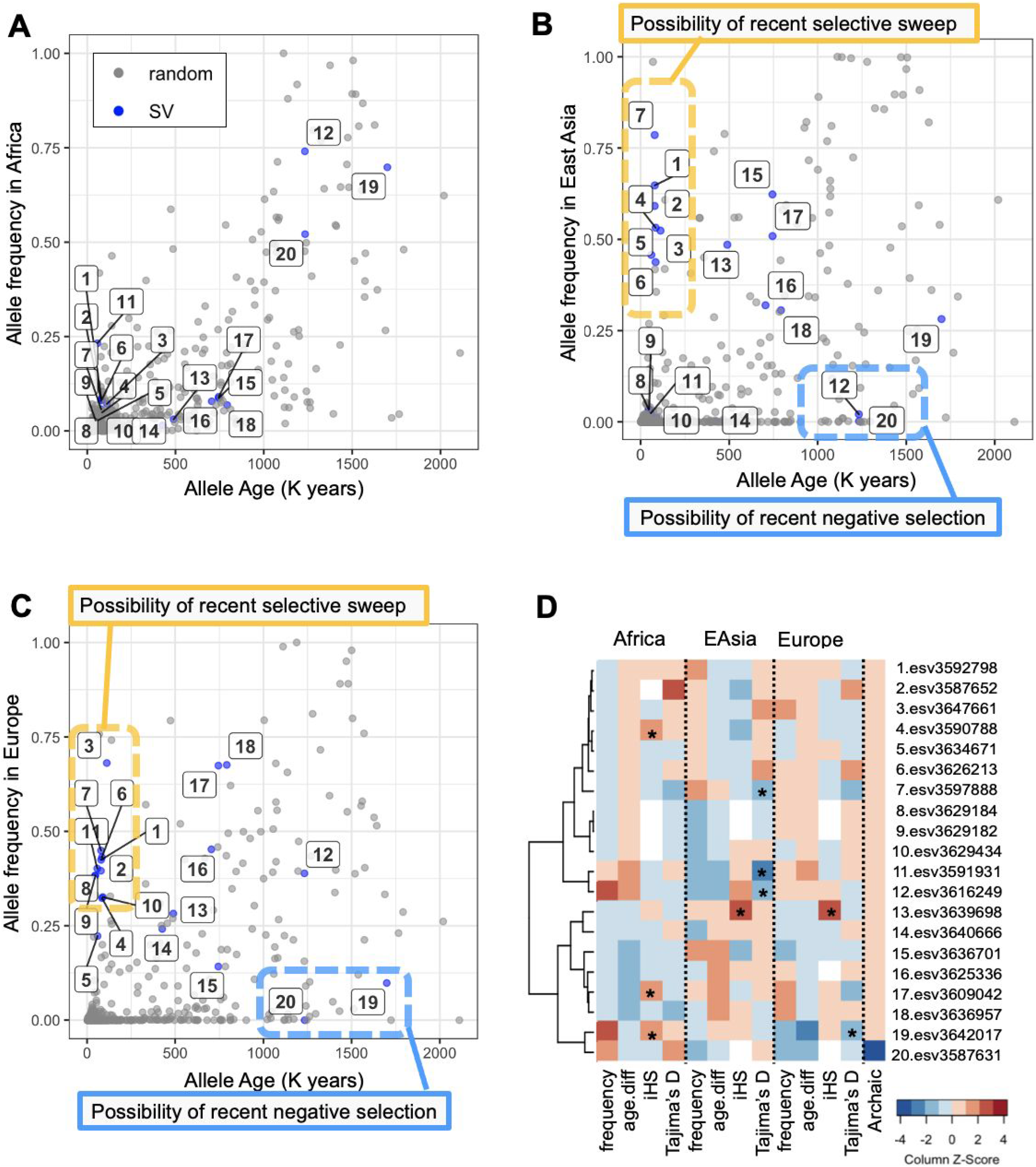
Neutrality tests on the 20 structural variants with phenotypic effects. **(A-C)** Age and frequency of the structural variants in African **(A)**, East Asian **(B)**, and European **(C)** populations. The variant’s ID is the same as **Table 2**. The blue dots represent the tag SNPs associated with the 20 structural variants (SV). The gray dots represent 599 random SNPs. The horizontal axis shows the age of allele in the Human Genome Dating database; the vertical axis is the alternative allele frequency in each population. **D.** A heatmap summarizing neutrality tests of the 20 structural variants in each population. It shows allele frequency, the allele age difference between haplotype-based estimation in the Human Genome Dating database (Albers and McVean 2020) and neutral expectation based on the allele frequency (Kimura and Ohta 1973), iHS value, Tajima’s D, and if the allele is observed in the Denisovan genome. (We did not observe these variants in the Neanderthal genomes.) Warmer colors indicate higher values. Similarly, for the allele age difference, colder and warmer colors show that the allele is older and newer than neutral expectation, respectively. For the archaic genome, blue shows that the allele is shared with the Denisovan. The asterisk indicates that the value is less than 5 percentile or more than 95 percentile when compared to at least 500 uniformly randomly selected variants or windows across the genome. (See Methods for detail.)

It is important to note that there are some limitations to our study. The most important is the samples that we use in our study is not a good representation of human variation overall. Further, these samples were not collected with specific adaptive pressures in mind (Scheinfeldt and Tishkoff 2013; Rees et al. 2020) Thus, our findings should be regarded as a methodological advance rather than a true snapshot of recent human evolution. Furthermore, for more detailed analyses, we focused on a subset of structural variants that are biallelic and present linkage disequilibrium with flanking variants, while hundreds of high-*MU* structural variants were multiallelic or without reported phenotypic effects in the GWAS ATLAS. These variants, which are associated with GWAS traits, tend to have higher allele frequencies in European populations. This is because the majority of GWAS have been conducted on European individuals (Sirugo et al. 2019), which is a side effect of the ascertainment bias in GWAS studies themselves. Nevertheless, these 20 structural variants and their associated haplotypes provide us with a glimpse into their evolution.

We found that the flanking haplotypes of three (esv3591931, esv3597888, esv3616249) and one (esv3642017) variants that reside in haplotype blocks show unusually low Tajima’s D values in East Asia and Europe, respectively (< 5th percentile compared to the genome-wide observations, **fig. 4D**). We found three variants (esv3642017, esv3609042, esv3590788) that tag single nucleotide variants with larger iHS values than the expected values (> 95th percentile compared to the genome-wide observations) in Africa. Another variant, esv3639698, is in linkage disequilibrium with single nucleotide variants that show unusually high iHS in both European and East Asian populations (> 95th percentile compared to the genome-wide observations). However, there is no consistent trend among these 20 haplotype blocks with unusually high *MU* values. For example, the haplotypes that show low Tajima’s D are not the same as the haplotypes that harbor SNPs with high XP-EHH or iHS values. Similarly, some of the haplotypes that show signatures of selection harbor structural variants that are extremely high in frequency (e.g., esv3642017 in Africa), while others harbor structural variants that are rare (e.g., esv3590788). Overall, our observations fit the emerging consensus in evolutionary genomics that the adaptive structural variants are shaped by complex evolutionary trajectories that change over time and space (Radke and Lee 2015; Mérot et al. 2020; Saitou and Gokcumen 2020).

As an example of the complicated nature of the evolutionary histories of adaptive structural variants, we highlight esv3642017. This variant is recorded as a deletion compared to the reference genome in the 1000 Genomes Phase 3 data set. However, a closer inspection reveals that this variant is a human-specific retro-insertion of the *DHFR* gene (Anagnou et al. 1988; Conrad et al. 2010; Schrider et al. 2013). The haplotype that harbors this insertion is associated with decreased height (p < 10^-16^). Even though deletion seems to be predominantly found in Africa, the derived retrogene inserted is predominantly found in Eurasia. The locus that harbors the insertion shows unusually low Tajima’s D in the European population (**fig. 4D**) and unusually low genetic diversity in another European-ancestry cohort as reported in Schrider et al. (2013), which altogether suggest a Eurasian-specific sweep of a recent insertion. Based on such locus-specific analyses, we identified incomplete population-specific sweeps and recent population-specific negative selection as the two main drivers for shaping the allele frequency distribution of putatively adaptive structural variants.

### Incomplete, Population-Specific Sweeps: The Intronic Deletion in Propionyl-CoA Carboxylase (PCC) Gene as a Case

The classical scenario for population-specific adaptive evolution is characterized by (i) high frequency of the variant in the specific population compared with other populations, (ii) deviations in the site frequency spectrum suggesting rapid expansion of the selected allele, resulting in an excess of rare variants in the locus, (iii) lower than expected allele age, and (iv) long haplotype homozygosity suggesting rapid expansion of the selected allele (Rees et al. 2020). We look for signatures of this scenario among the 20 haplotypes that we highlight because they harbor structural variants with unusual allele frequency distributions (*MU* in the 1st percentile, > 4.23) and because they are associated with GWAS traits (**Table 2**). We found that 12 (60%) of them fit the scenario of recent population-specific adaptive sweep (**fig. 4A-D**).

The haplotype harboring esv3597888 provides an informative example of the population-specific incomplete sweep scenario. The haplotype has a lower than 5% allele frequency in most African populations but reaches near 75% allele frequency in East Asian populations (**fig. 5A**). Further, the Median Joining network of the haplotypic variation in this locus shows a dramatic reduction of haplotypic diversity beyond the expected reduction due to drift in East Asian population as compared with African population, which is consistent with a recent selective sweep (**fig. 5B**). The Tajima’s D values retrieved from the flanking sequences of the deletion are lower than genome-wide expectations in all three continental populations (**fig. 5C**). Last but not least, the estimated age of the allele is much more recent than what is expected based on its frequency, especially in the East Asian population (**fig. 4B, fig. S3**). Collectively, these results suggest a recent selective sweep in Eurasian populations. However, even for this locus, not all the neutrality tests capture this sweep. For example, in a traditionally defined recent sweep, we expect to find high iHS values. Instead, for this locus, the iHS is relatively low, mirroring the surprisingly high overall haplotypic variation in this locus (**fig. 4C**). Regardless, esv3597888 remains one of the best candidates for a derived structural variant that has recently been swept to higher allele frequency in a population-specific manner.

**Figure 5:**
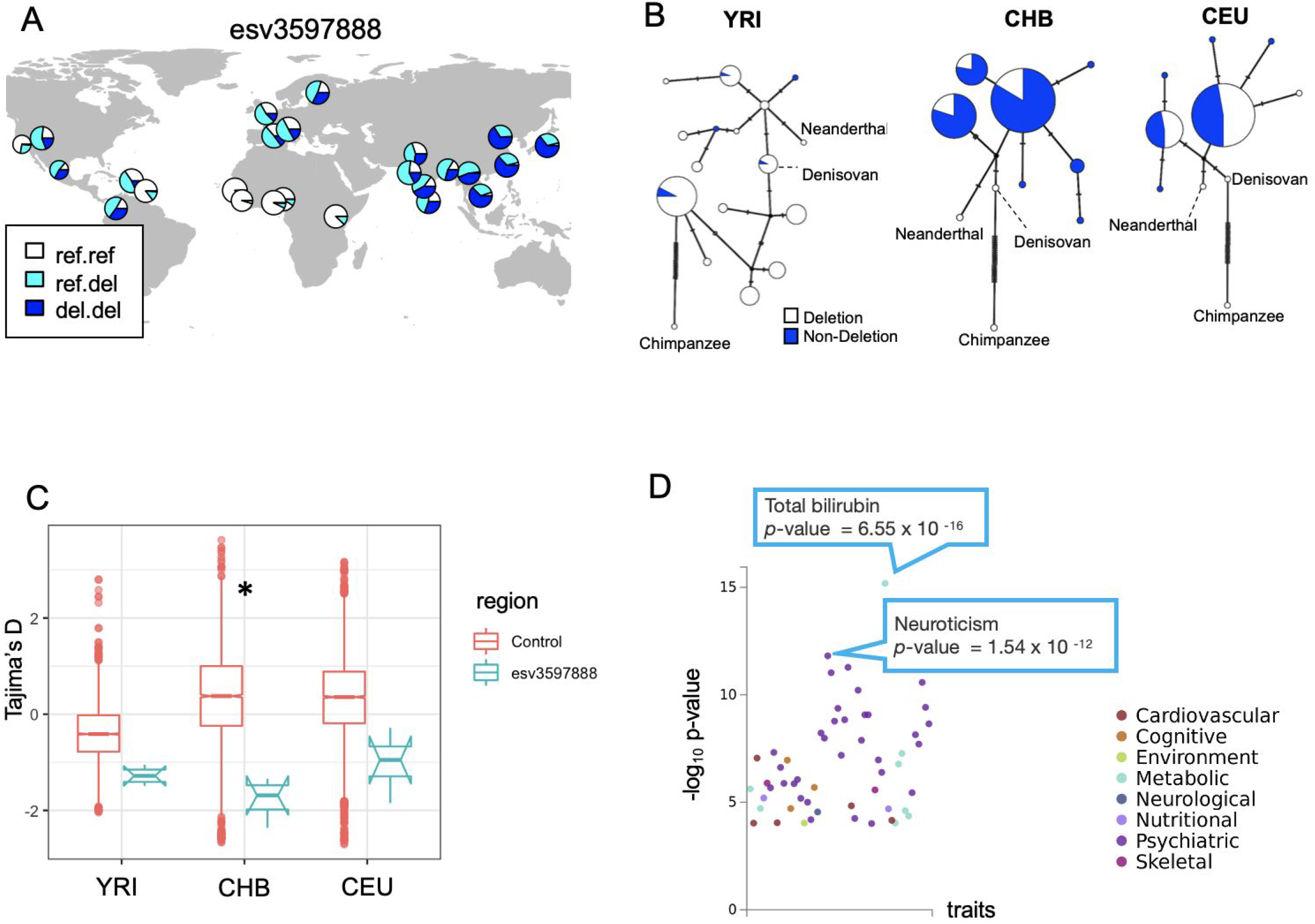
The evolutionary analysis of esv3597888, the deletion overlapping the intronic region of the propionyl-CoA carboxylase (PCC) gene. **A.** The geographic distribution of the esv3597888. **B.** Haplotype networks constructed from the 5kb upstream and downstream sequences from the esv3597888 location of three modern human populations (Yoruban (YRI), Han Chinese (CHB), and European (CEU),), the Altai Neanderthal sequence (Prüfer et al. 2014) and the Denisovan sequence (Reich et al. 2010) that are mapped to hg19 reference genome, and the chimpanzee reference genome (panTro4). The haplotypes that harbor the deletion are indicated by white and those that do not are indicated by blue. **C.** Tajima’s D value in the 5kb upstream and downstream regions of esv3597888 (10kb in total) (Tajima 1993). Asterisk shows that Tajima’s D of the esv3597888 flanking region is lower than the bottom five percentile of Tajima’s D of 5000 random regions. The asterisk shows that the mean value of esv3597888 tag region is lower than the five percentile of the control region. **D**. The PheWAS result of rs556788, which tags esv3597888 (**Table S4**). Each dot indicates a trait. The vertical axis shows the -log10 p-value of the association between the genotype and phenotype. Color indicates the phenotype category in GWAS ATLAS.

**Figure 6:**
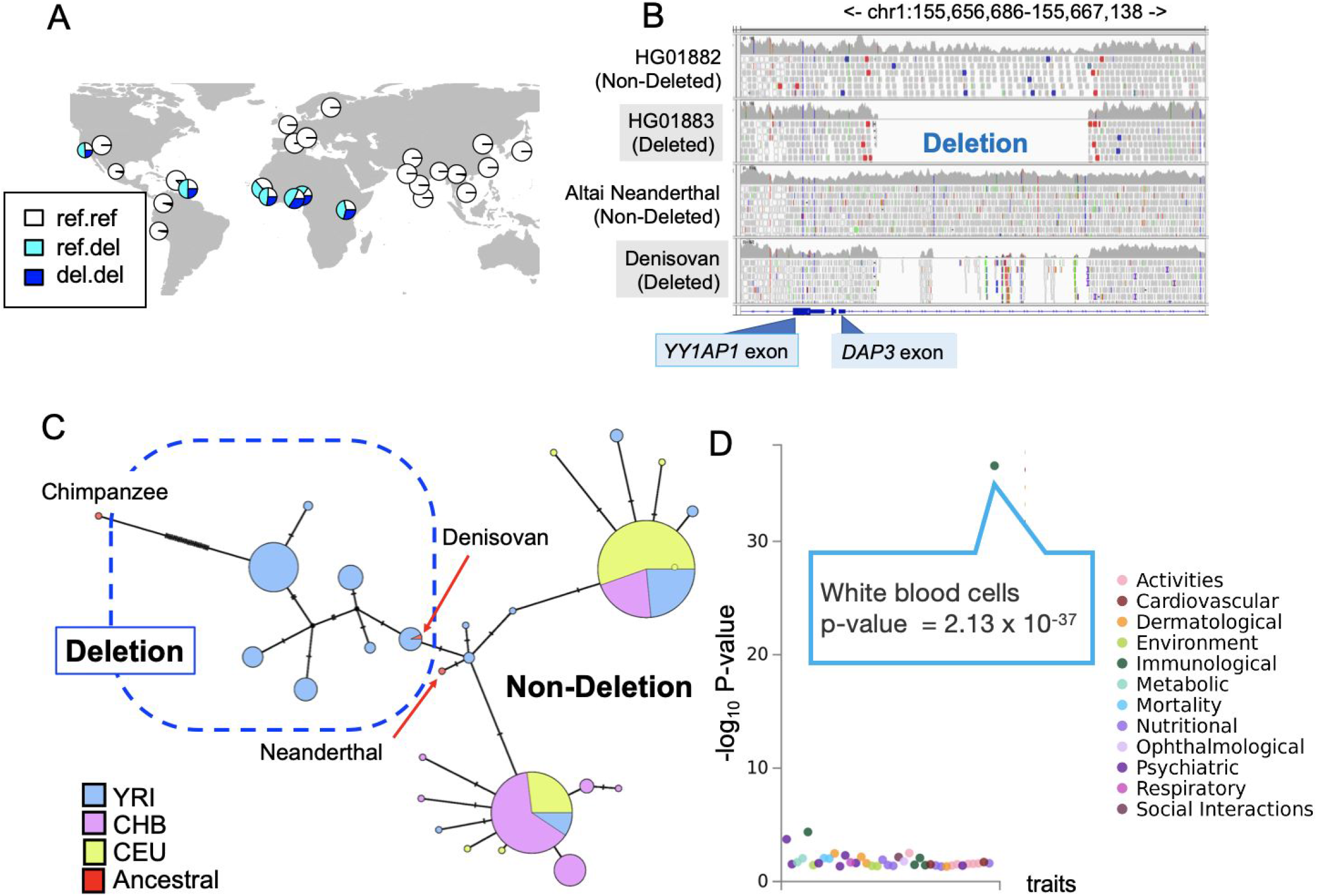
The evolutionary analysis of esv3587631, the intronic deletion polymorphism of the death-associated protein gene3 (DAP3). **A.** The geographic distribution of the esv3587631. **B**. esv3587631 in modern and ancient hominin genomes. These Integrated Genome Browser snapshots show the genome assembly (Hg19) of a human with the ancestral, homozygous non-deleted genotype and another with a homozygous deleted genotype that shows no reads mapping to the deletion region (top two rows). Similarly, sequences from a Neanderthal and a Denisovan genome assemblies were mapped to this region. The Denisovan genome shows a clear signature of the deletion with breakpoints indistinguishable from the deletion observed in modern humans. **C.** A haplotype network of three modern human populations, Yoruba (YRI), Han Chinese (CHB), European (CEU), as well as Altai Neanderthal and the Denisovan, and Chimpanzee, constructed from the flanking sequences from the esv3587631 location. **D.** The Phewas result of rs348195, the tag SNP of esv3587631 at GWAS Atlas. Each dot indicates a trait and the y-axis shows the -log_10_ p-value of the association between the genotype and phenotype.

The phenotypic effects of the haplotype harboring this variant further support the potential adaptive relevance of esv3597888. This 5.4kb deletion overlaps with the intronic region of the propionyl-CoA carboxylase (PCC) gene, which encodes for an enzyme that metabolizes specific amino acids and lipid species (Wongkittichote et al. 2017). The haplotype harboring esv3597888 (tagged by rs556788) is associated with the expression of the PCCB gene in the adrenal gland (p = 7.8 × 10^−10^ in the GTEx Analysis Release V8 (GTEx Consortium 2013)). Moreover, the haplotype is associated with total bilirubin, a cardiometabolic signaling molecule (p = 6.6 × 10^−16^), and neuroticism (p = 1.5 × 10^−12^) (**fig. 5D**). We argue that it is likely that the 5.4kb deletion esv3597888 is the causal variant in these associations, given its size and that it overlaps with a well documented binding site for the abundant transcription factor CTCF (**fig. S4**). The haplotype has pleiotropic functional effects, and thus the exact reasons why it confers an adaptive advantage in East Asia particularly remains to be seen.

### Recent Population-Specific Negative Selection: Ancient Intronic Deletion in Death-Associated Protein (DAP3) Gene as a Case

Among the haplotypes that we highlighted, we noticed that some show unexpectedly low allele frequency in Eurasian populations compared to the expectation based on their estimated age (**Table 3**, **fig. 3B and 3C**). We hypothesize that these haplotypes, and by proxy the structural variants that they harbor, may have been subjected to recent, population-specific negative selection, favoring the ancestral allele. Variations that have emerged early in human evolution and remain in extant populations are often found at high allele frequencies in all extant human populations under neutrality (Lin et al. 2015). Thus, if *recent* negative selection on the derived structural variant is acting in a population-specific manner, we expect to observe in that population i) an unusual reduction in allele frequency of the variant that cannot be explained by drift alone and ii) a shift in the allele frequency spectrum towards rare variants in the locus.

A striking example of population-specific negative selection is provided by the haplotypes harboring esv3587631, which shows one of the highest *MU* values in the genome (*MU* = 8.50). This deletion is the major allele (i.e., > 50% allele frequency) in most sub-Saharan African populations but almost absent in non-African populations (**fig. 4A**). Human Genome Dating database estimates the age of a single nucleotide variant tagging esv3587631 to be 1,1-1,3 Million years old (Albers and McVean 2020). Thus, the deletion has emerged prior to human-Neanderthal divergence. Consistent with this result, we found that Denisovan but not Altai Neanderthal carries this deletion (**fig. 4B**). The haplotype network showed that the haplotypes harboring the deletion are similar to those from archaic hominins, consistent with our observation that this deletion is present in archaic human genomes (**fig. 4C**). Collectively, it is clear that the deletion has evolved before Human Neanderthal divergence and increased in allele frequency in African populations to more than 75%, harbored by diverse haplotypes. However, none of the haplotypes that harbor the deletion is found in Eurasian populations. Furthermore, the locus shows significantly negative Tajima’s D values in the East Asian population, further supporting non-neutral forces acting on the deletion (**fig. 5D**).

Functionally, this ~4.8kb deletion overlaps with one of the introns of the well-studied and highly conserved *DAP3* gene (**fig. 4B**). DAP3 is a mitoribosome protein that regulates apoptosis at the cellular level and is linked to multiple developmental, immune-related, and biomedically relevant phenotypes at the organismal level (Greber and Ban 2016; Kim et al. 2017). Specifically, the deletion overlaps with a conserved regulatory region comprising multiple transcription factor binding sites (**fig. S4**). Consistent with these observations, the haplotypes harboring the deletion (tag variant, rs348195) was strongly associated with the increased expression of the *DAP3* gene in various tissues (p < 10^−6^), with the effect size exceeding 0.4 in some cases. Moreover, the deletion (through the analysis of tag SNP rs348195) is strongly associated with decreased levels of white blood cells (p = 2.1 × 10^−37^, **Table 2**). Collectively, these results are consistent with a scenario where an ancient deletion variant that has been either neutral or beneficial in African populations has become detrimental to fitness in Eurasian populations, perhaps due to adaptive constraints concerning immune function.

## Conclusion

Although several putatively adaptive structural variants have been reported in previous studies, a genome-wide selection scan of structural variants has remained challenging. In this study, we built a network-based analysis of population differentiation among 26 populations in the 1000 Genome Project data set to identify putatively adaptive structural variants including multi-allelic variants. Our method assumes that drift is the major force that shapes the distributions of genomic variants among human populations as articulated by others (Ramachandran et al. 2005; Coop et al. 2009). In identifying the most common allele frequency distribution combinations across the 26 populations, our method parallels the recent variant-centric integrative analysis method proposed by Biddanda et al (2020). We argue that such direct, empirical scrutiny of the geographical distribution of variants will provide a valuable and relatively unbiased picture of demographic and non-neutral trends that shape human genetic variation.

Our method is designed to identify structural variants with population differentiation that deviate from neutral expectations without any *a priori* hypothesis. It identified hundreds of putatively adaptive structural variants with unusual genotype frequency distributions in humans. The majority of these structural variants were hidden from traditional selection scans which mainly focus only on bi-allelic single nucleotide variants. Our results include 24 exonic multiallelic structural variants, the majority of which were not discussed within an adaptive context in humans. In addition to incomplete sweeps of derived structural variants, we found that recent population-specific negative selection is a considerable force shaping the geographic distribution of functional structural variants in humans. Overall, our study supports the emerging notion that structural variants significantly contribute to non-neutral and biomedically relevant phenotypic variation in humans (Radke and Lee 2015; Mérot et al. 2020) and highlight specific trajectories underlying the evolution of such variants.

From an evolutionary genomics perspective, the prominence of exonic multiallelic copy number variants among the putatively adaptive structural variants is not surprising. Cross-species analyses have repeatedly revealed the outsized role of recurrent gain and losses in gene families in shaping phenotypic characteristics in a variety of species, with recurrent evolution of caffeine in plants (Denoeud et al. 2014), salivary amylase in mammals (Pajic et al. 2019), and venom in snakes (Casewell et al. 2020) providing notable examples. Moreover, studies in humans reported that multiallelic copy number variants have seven times more effect on gene dosage than the combined effect of biallelic deletions and duplications (Handsaker et al. 2015). The same multiallelic structural variants, however, are hidden in the majority of GWAS and selection analyses. Multiallelic variants are not necessarily tagged by nearby single nucleotide variants, and they often reside in the genomic regions with enriched segmental duplications where identifying variants can be problematic. Thus, we expect that better genotyping of multiallelic structural variants with long-read sequencing platforms will dramatically increase our ability to identify multiallelic structural variants and their previously unknown adaptive roles.

A surprising result from our study is the identification of recent negative selection favoring ancestral alleles as a notable force determining the allele frequency distribution of putatively adaptive structural variants. Selective sweeps are often thought to increase the allele frequency of the *derived* and not *ancestral* variant. In this work, we found that at least 10% of the putative adaptive structural variants show recent sweeps favoring the ancestral allele. It is plausible that recent human adaptive evolution involves repeated adaptation to similar environmental conditions across time and geography as reported in (Bergey et al. 2018). Thus, an ancestral adaptive variant that confers a smaller fitness advantage than the derived variant may become adaptively beneficial again if environmental pressures revert back to an earlier state. This scenario is particularly applicable to immune-system-related traits within the context of an evolutionary arms race as articulated previously (Key et al. 2014). Similarly, adaptive landscapes concerning metabolic traits have drastically changed multiple times for human populations due to technological advances (e.g., agricultural transition) (Hancock et al. 2010) and migrations to new ecologies (e.g., arctic populations) (Marciniak and Perry 2017). Thus, under the assumption that neither the ancestral or derived alleles are fixed, it is not surprising that ancestral structural variants are favored in certain geographies and instances. Such cases will appear as negative selection against the derived allele. We reported in detail one such case involving the exonic deletion of the growth hormone receptor in another study (Resendez et al.). The current study identifies several other cases, suggesting that recent, geography-specific negative selection is a considerable force shaping allele frequency distribution and population differentiation of functional structural variants.

There are caveats to our study and to the investigation of adaptive structural variants in general. First, it is clear from existing literature that the current data sets suffer from significant false-positive rates, potentially missing up to 80% of the structural variants (Mahmoud et al. 2019). Moreover, current technologies can discover certain types of structural variants (e.g., large biallelic deletions) much more sensitively than other types of variants (e.g., duplications, inversions). It is telling that one of the structural variants most relevant to human evolution, amylase copy number variation, are not cataloged by the 1000 Genomes Phase 3 data set because of alignment issues in the locus. Even when such multiallelic variants are discovered, it is not uncommon that their exact genotypes (e.g., exact copy number) may not be accurately documented. Second, the genotyping platforms commonly used in genetic association studies mostly focus on bi-allelic single nucleotide variants only. In fact, even this study, which is aware of these limitations, highlighted bi-allelic variants, for which the haplotype can be readily resolved, and thus trait associations can be investigated. The true contribution of most structural variants, including multiallelic variants, to phenotypic variation, remains mostly unknown. Overall, the current picture of the evolutionary impact of structural variants, including that revealed in this study, remains incomplete and should be treated as a theoretical and methodological framework for future studies with more comprehensive data sets. We believe that as long-read sequencing-based discovery and later genotyping become affordable, the full impact of structural variants on human evolution and diversity will be better revealed.

## Methods

### 1000 Genomes Phase 3 data set

As the input data set, we used 1000 Genome Project phase 3 data sets (Sudmant, Rausch, et al. 2015) for the following three reasons. First, the genotyping is based on whole-genome sequencing and multiple detection methods such as Delly (Rausch et al. 2012), which combines short insert paired-ends, long-range mate-pairs, and split-read alignments, and GenomeSTRiP (Handsaker et al. 2011), which uses read-depth and read pairs for structural variant identification to improve accuracy. Thus, this data set provides a highly accurate structural variant genotype. Second, it contains approximately 100 individuals from each population. Therefore, one can increase the power to detect geographically-differentiated structural variants due to population-specific adaptation by assessing deviations from expected population differentiation. Third, it provides phased genotype information not only of the structural variants but also of the SNPs from the same individuals. This allows us to apply our methods for identifying population differentiation to known SNPs to assess their performance and to carry out the subsequent haplotype-based analysis on a subset of structural variants.

### Pre-processing structural variants and selection of known SNPs in the analysis

We selected 57,629 autosomal structural variants with annotations in the 1000 Genomes project phase 3 data set (Sudmant, Rausch, et al. 2015) since variants in sex-chromosomes are differently described from autosomes due to the smaller number of the observed number of chromosomes and can not be analyzed in the same pipeline as autosomal variants. As controls, we also used 1008 uniformly randomly selected single nucleotide variants from the same data set and six single nucleotide variants that have undergone putative natural selection, including rs334 in HBB (Ding et al. 2013), rs73885319 in APOL (Ko et al. 2013), rs4988235 in LCT(Smith et al. 2009), rs12913832 in OCA (Wilde et al. 2014), rs3827760 in *EDAR*, which is common in East Asian populations and associated with hair and dental traits (Mou et al. 2008; Kimura et al. 2009; Wu et al. 2016), rs1426654 in SLC24A5, which is associated with skin color (Norton et al. 2007; Basu Mallick et al. 2013; Deng and Xu 2018). In addition to these SNPs we included rs1129740 that falls into *HLA-DQA1*, which is one of the few variants in the human genome that showed classical signatures of balancing selection (Teixeira et al. 2015). This *HLA* allele showed unusually low *MU* (0.22) despite the global allele frequency of 0.52 (**fig. 2**).

### Calculation of the MU and MDS plot

We characterize each locus *ℓ* as an *N* ✕ *N* similarity matrix, denoted by *S_ℓ_*, where *N* = 26 is the number of populations in the 1000 Genome Project Phase 3 data set (**fig. 1A**). The entries of matrix *S_ℓ_* represent the similarity between pairs of populations in terms of the frequency of each allele at a locus. Specifically, for each locus *ℓ* and population *i*, the genotype count is given by the 1000 Genome Project Phase 3 data set. In general variant call format (VCF), genotype (i.e., the allele status of a pair of chromosomes of one individual) is denoted by *x*_1_|*x*_2_. Genotypes 1|0 and 0|1 in general variant call format are effectively the same and mean that one individual has one reference (i.e., 0) allele and one alternative (i.e., 1) allele at the locus. Therefore, we summarized both 1|0 and 0|1 into 0|1 in the following analysis. The maximum (alternative + reference) allele number at a locus was nine, in which case the allele number ranges from 0 to 8. Therefore, in general, we summarized *x*_1_|*x*_2_ and *x*_2_*|x*_1_ into *x*_1_|*x*_2_, where *x*_1_, *x*_2_ = 0,1, …, 8 and *x*_1_ ≤ *x*_2_. We denote the frequency of genotype *x*_1_|*x*_2_ at locus *ℓ* and population *i* by *p_ℓ_*_,*i*_ (*x*_1_|*x*_2_), where 0 ≤ *x*_1_ *≤ x*_2_ *≤* 8. Note that 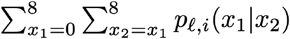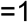. To increase the sensitivity to identify the population differentiation of multiallelic variants (i.e., variants with more than two alleles), especially, with large copy number variation (such as a multiallelic variant with copy number one to eight, even if the frequency of copy number eight is rare), we use a weighted variant of the Bhattacharyya similarity metric, which modifies the Bhattacharyya similarity metric (Bhattacharyya 1943; Cha and Srihari 2002), as follows (**fig. 1B**).

First, we transform the original distribution,{p_ℓ_,_i_ (*x*_1_ | *x*_2_;) 0 ≤ *x*_1_ ≤ *x*_2_ ≤ 8} to 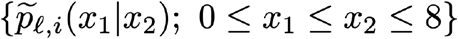
, where

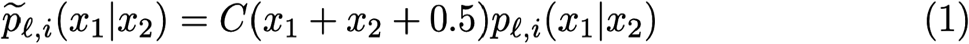

and

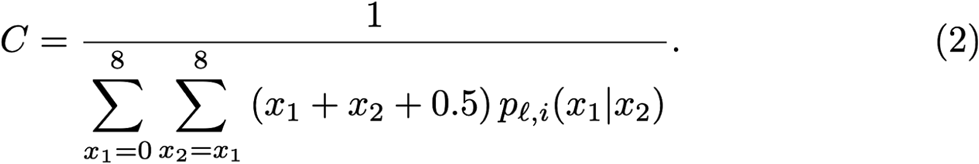

This transformation magnifies the frequency of genotype and its differences between populations at large *x*_1_ and *x*_2_ values (i.e., large copy number variation). Second, we measure the Bhattacharyya metric between each pair of populations *i* and *j* based on the transformed probability distribution, *i.e.*,

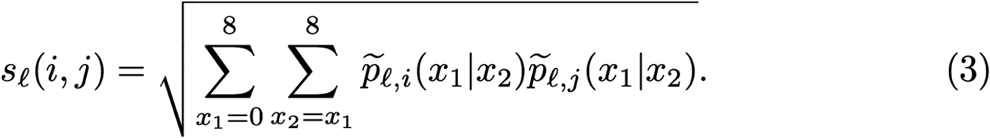

If the two distributions 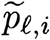 and 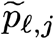 are identical, then *s_ℓ_* (*i, j*) = 1, which is the largest value of *s_ℓ_* (*i, j*). In this manner, for each locus *ℓ*, we obtain an *N* ✕ *N* similarity matrix *S_ℓ_* whose (*i, j*) entry is given by Equation (1). To analyze the organization of *M* = 58,644 loci (which is composed of 57,629 structural variants, 1,008 random single nucleotide polymorphisms, and seven single nucleotide polymorphisms under adaptive evolution), we constructed a distance matrix for the *M* loci by comparing similarity matrix *S_ℓ_* across the loci. We define the distance between *S_ℓ_* and *S_ℓ′_* using the Frobenius norm, denoted by *d*_F_, which is given by

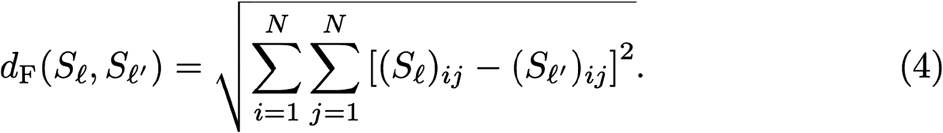

Note that (*S_ℓ_*)_i*j*_ = s_*ℓ*_(*i*, *j*). Also note that one can replace the summation by 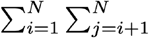 without loss of generality because the similarity matrix *S_ℓ_* is symmetric and all of its diagonal elements are equal to 1. The *M* ✕ *M* Frobenius distance matrix, denoted by *F*, tabulates the difference between each pair of loci, and its (*ℓ*, *ℓ′*) entry is given by *d*_F_(*S_ℓ_*,*S_ℓ_*_′_). Finally, we ran a multidimensional scaling (MDS) algorithm on *F* to map out relationships between the *M* loci on a two-dimensional space. We used the Python package *manifold*, which is part of *scikit-learn* (Pedregosa et al. 2011), to estimate the MDS. We only used one initial condition due to the long time required for the computation. The analyses have been done on the server of the University at Buffalo Center for Computational Research (http://www.buffalo.edu/ccr.html).

### Allele frequency, *MU* and structural variants

Since *MU* is related to alternative allele frequency by definition, we categorized the variants into four groups in terms of the allele frequency using ranges 1-0.25, 0.25-0.5, 0.5-0.75, and 0.75-1, and investigated the distribution of *MU* of the variants in each group. In intermediate allele frequency groups, multiallelic copy number variants (CNV) had higher *MU* than SNPs (Wilcoxon test, p = 0.0021 and p = 0.00095 for allele frequency ranges 0.25-0.5 and 0.5-0.75, respectively). This result indicates that our methods may detect population differentiation due to the excess of multi-copy alleles than biallelic variants.

### Functional genomics analysis

We retrieved exonic content from UCSC Genome Browser (http://genome.ucsc.edu/; UCSC Genes, table: knownCanonical) and examined if each structural variant contained one or more entire exon using bedtools (Quinlan and Hall 2010). To find functionally relevant loci, we first calculated linkage disequilibrium between the top 1% high *MU* biallelic structural variants and neighboring regions (5kb upstream and downstream) with vcftools (Danecek et al. 2011). We searched the resulting tag SNPs (r^2^ > 0.95) in GWAS Atlas Phewas database (https://atlas.ctglab.nl/PheWAS), defining that p < 10^−9^ as a statistically significant association. Of the 576 top 1% structural variants in terms of *MU*, we found that 500 variants were bi-allelic, which were suitable for haplotype analysis. Among the 500 variants, 344 structural variants showed R^2^ larger than 0.95 with neighboring variant(s). Of these 344 structural variants, 20 of them have flanking tag SNPs that are significantly associated with phenotypic variation (**fig. 2A, Table 2**). Further, we used the GTEx portal (GTEx Consortium 2013) for associating these SNPs to variation in gene expression levels (fig. S4).

### Evolutionary Genomics Analysis of the haplotypes harboring putatively adaptive structural variants

We used the 344 structural variants that are among the top 1% in terms of *MU* and have flanking tag SNPs with R^2^ value larger than 0.95 for the following evolutionary analysis. We used the age estimates from Human Genome Dating database (Albers and McVean 2020) using tag SNPs as proxies to the adaptive SVs. This database documents the allele age estimates based on the analysis of pairwise haplotype identical tracts in 1000 Genomes (The 1000 Genomes Project Consortium et al. 2015) and Simons Genome Diversity Projects (Brazma 2019) (https://human.genome.dating; last accessed, 3.23.2020). We also calculated how long it takes for a new allele to reach a given frequency in each population under neutrality using equation (15) of (Kimura and Ohta 1973). For this calculation, we used demographic models for each population detailed in (Schaffner et al. 2005) (**fig. 3D**).

For haplotype-level population genetics measures, we targeted 5k upstream and downstream regions of the structural variants and calculated Tajima’s D scores of the tag SNP of the target structural variants using VCFTools (Danecek et al. 2011). We retrieved the iHS score from the 1000 Genomes Selection Browser (Pybus et al. 2014). For comparative purposes, we calculated the same scores for ~500 random regions generated with bedtools (Quinlan and Hall 2010) across the genome. Ancient human (Altai Neanderthal and Denisovan) genomic bam files are published on the Max-Planck Institute website (https://www.eva.mpg.de/index.html) (Reich et al. 2010; Prüfer et al. 2014). We used samtool-based (Li et al. 2009) read-depth analysis to genotype deletions in archaic genomes (**fig. 5B**). We generated haplotype networks using VCFtoTree (Xu et al. 2017) and POPArt (Clement et al. 2002; Leigh and Bryant 2015).

## Supporting information

STables

## Data visualization

For the data visualization, we used Rstudio (v1.2.1335), R(v3.5.3), and ggplot2 (Wickham 2009). Codes will be available at https://github.com/GokcumenLab.

## Acknowledgement

We thank Drs. John Novembre and Simen Rød Sandve for the careful reading of this manuscript.

## SUPPLEMENTARY Materials

**Table S1** All the variants and *MU* values

**Table S2** Notable variants identified in this study (Low *MU* and High frequency; bottom 25% *MU* and 0.4 − 0.6 allele frequency,)

**Table S3** High *MU* (i.e., top 1%) exonic variants

**Table S4** All the variants with high *MU* (top 1%) and high LD (R^2^ > 0.95) with neighbouring variants

---

**Figure S1.**
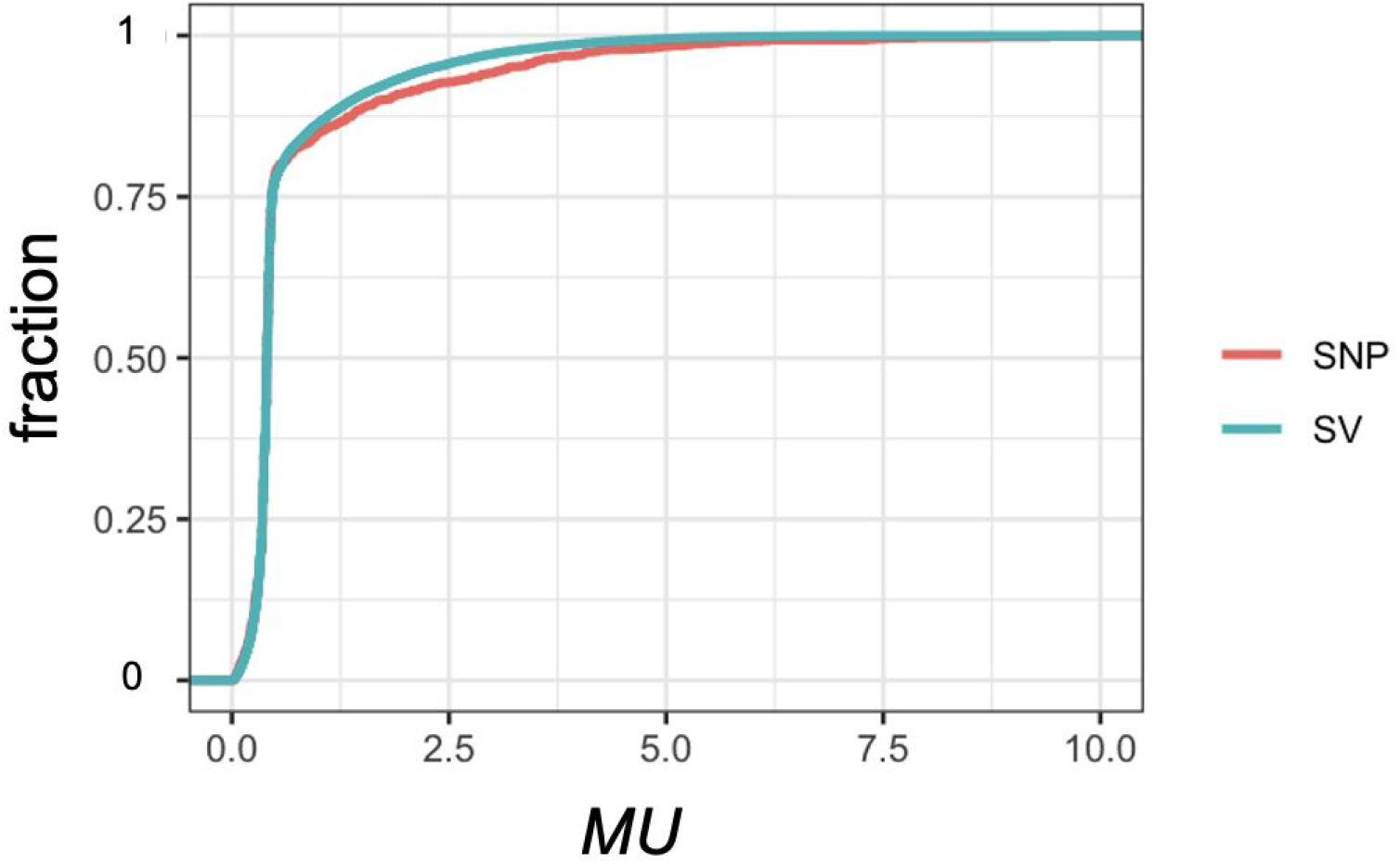
Cumulative fraction of the *MU* of SNPs and structural variants (SV)

**Figure S2.**
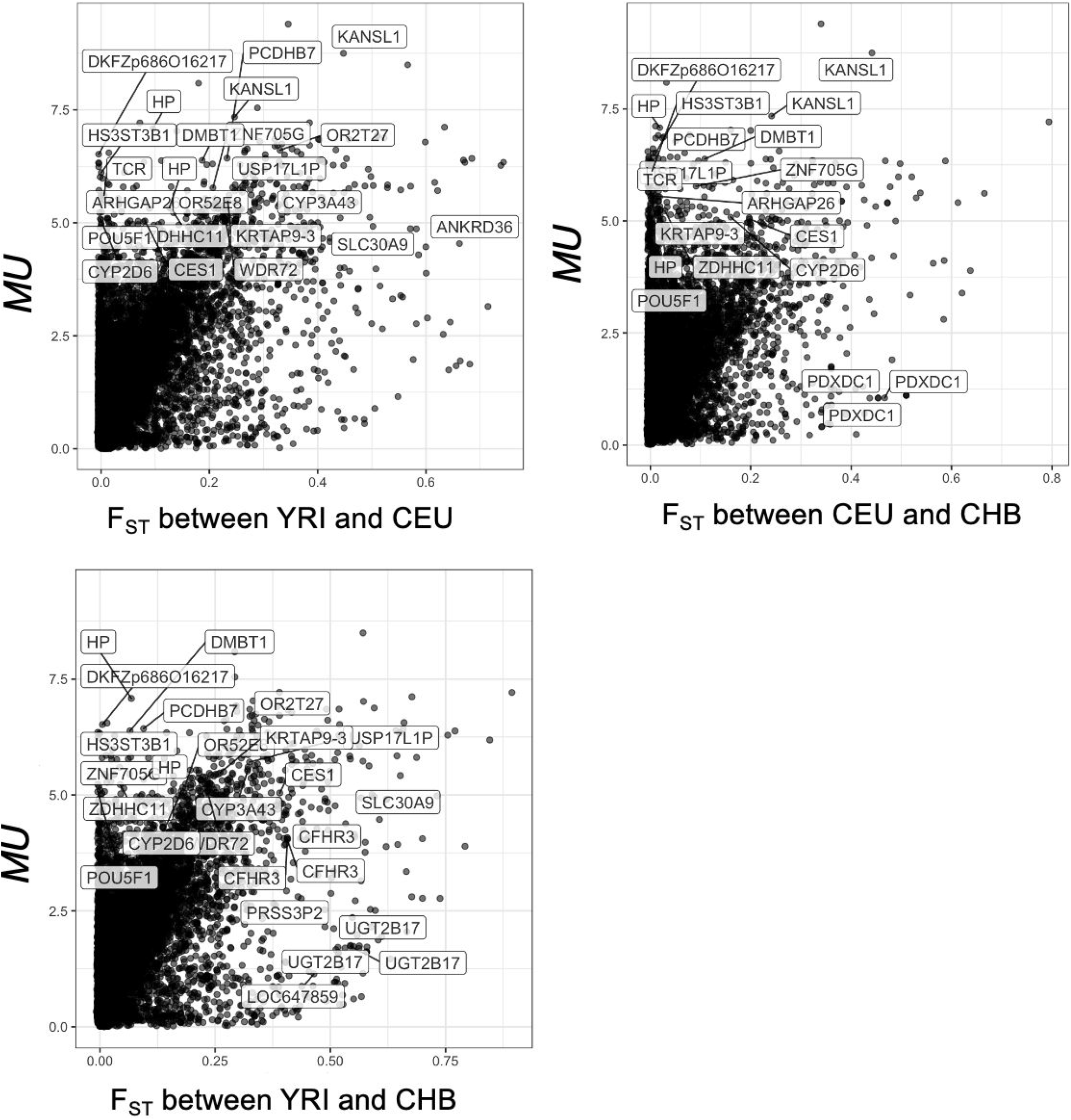
Comparison between F_ST_ and *MU* values of structural variants

**Figure S3.**
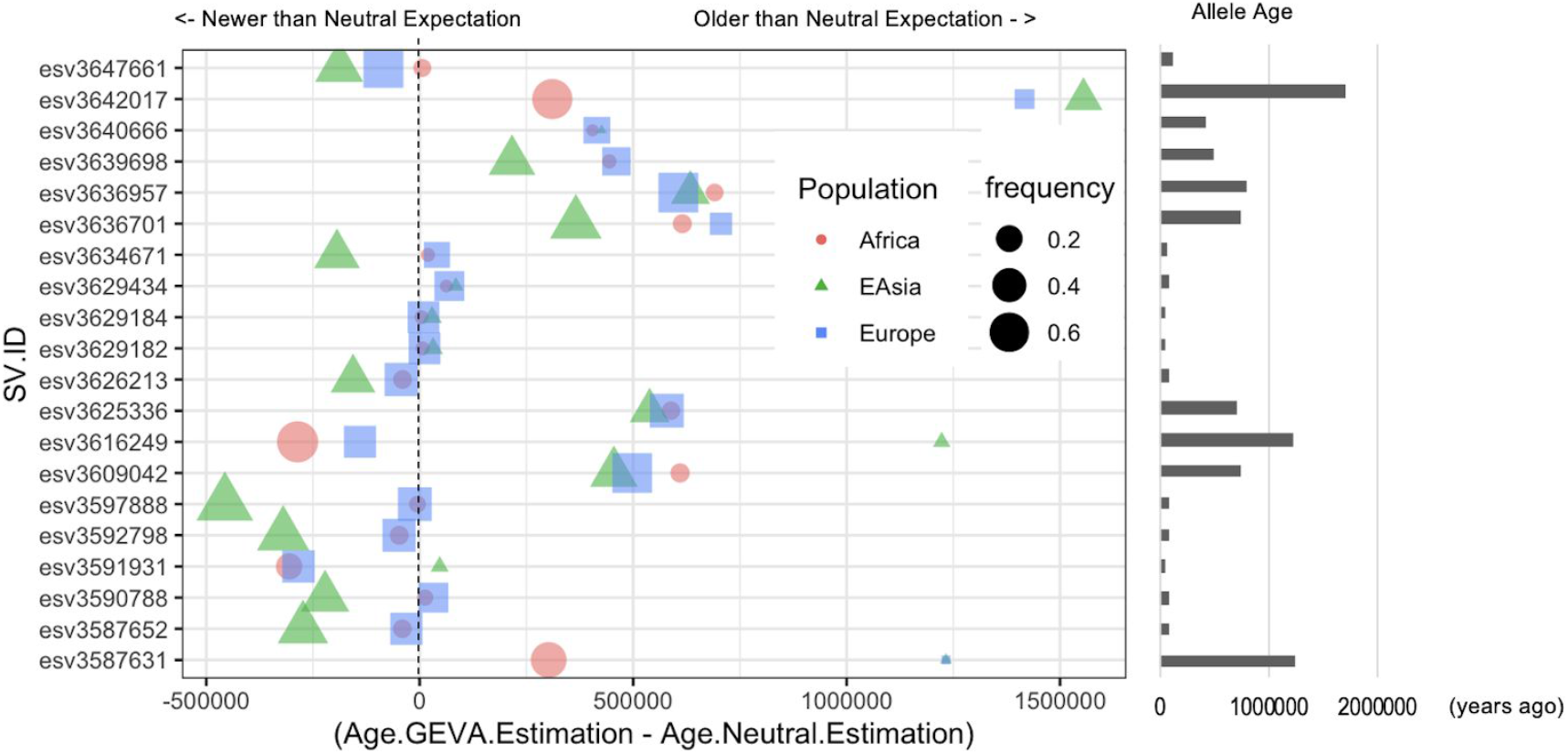
The difference between haplotype-based age estimation and expected age under neutral evolution for the 20 structural variants with phenotypic effects (**Table 2, Figure 4D**)

**Figure S4.**
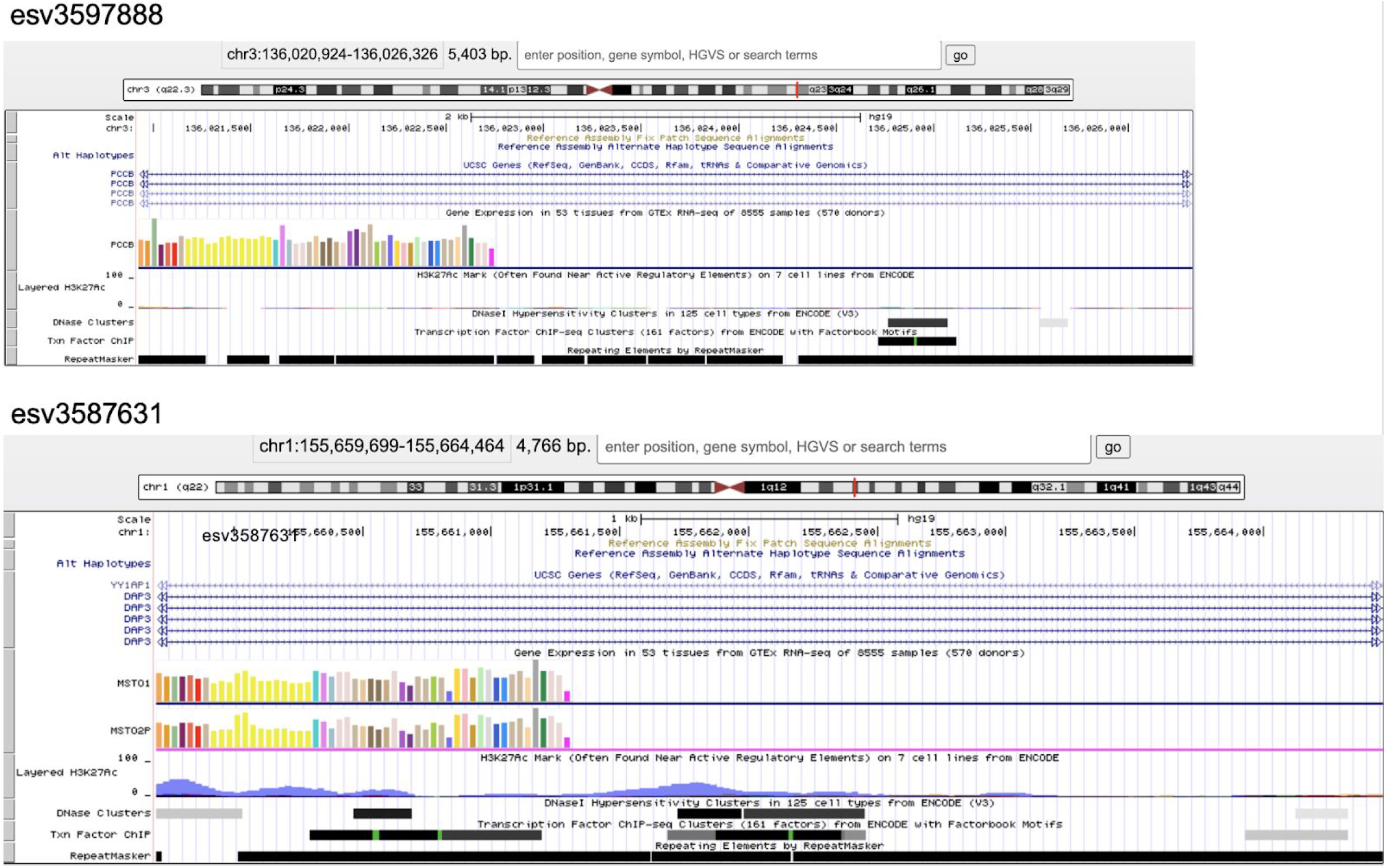
The overlap between functional annotation and structural variants

